# MG7: Configurable and scalable 16S metagenomics data analysis

**DOI:** 10.1101/027714

**Authors:** Alexey Alekhin, Evdokim Kovach, Marina Manrique, Pablo Pareja-Tobes, Eduardo Pareja, Raquel Tobes, Eduardo Pareja-Tobes

## Abstract

As part of the Cambrian explosion of omics data, metagenomics brings to the table a specific, defining trait: its social essence. The *meta* prefix exerts its influence, with multitudes manifesting themselves everywhere; from samples to data analysis, from actors involved to (present and future) applications. Of these dimensions, data analysis is where needs lay further from what current tools provide. Key features are, among others, scalability, reproducibility, data provenance and distribution, process identity and versioning. These are the goals guiding our work in MG7, a 16S metagenomics data analysis system. The basic principle is a new approach to data analysis, where configuration, processes, or data locations are static, type-checked and subject to the standard evolution of a well-maintained software project. Cloud computing, in its Amazon Web Services incarnation, when coupled with these ideas, produces a robust, safely configurable, scalable tool. Processes, data, machine behaviors and their dependencies are expressed using a set of libraries which bring as much as possible checking and validation to the type level, without sacrificing expressiveness. Together they form a toolkit for defining scalable cloud-based workflows composed of stateless computations, with a static reproducible specification of dependencies, behavior and wiring of all steps. The modeling of taxonomy data is done using Bio4j, where the new paradigm of graph databases allows for both a simple expression of taxonomic assignment tasks and the calculation of taxa abundance values considering the hierarchic structure of the taxonomy tree. MG7 includes a new 16S reference database, *16S-DB7*, built with a flexible and sustainable update system, and the possibility of project-driven personalization.

## Introduction

During the past decade, metagenomics data analysis is growing exponentially. Some of the reasons behind this are the increasing throughput of massively parallel sequencing technologies (with the derived decrease in sequencing costs), and the wide impact of metagenomics studies [1], especially in human health (diagnostics, treatments, drug response or prevention) [2]. We should also mention what could be called the microbiome explosion: all kind of microbiomes (gut, mouth, skin, urinary tract, airway, milk, bladder) are now routinely sequenced in different conditions of health and disease, or after different treatments. The impact of Metagenomics is also being felt in environmental sciences [3], crop sciences, the agrifood sector [4] and biotechnology in general [5] [6]. These new possibilities for exploring the diversity of micro-organisms in the most varied environments are opening new research areas, and drastically changing the existing ones.

As a consequence, the challenge is thus moving (as in other fields) from data acquisition to data analysis: the amount of data is expected to be overwhelming in a very short time [7].

Genome researchers have raised the alarm over big data in the past [8], but even a more serious challenge might be faced with the metagenomics boom. If we compare metagenomics data with other genomics data used in clinical genotyping we find a differential feature: the key role of time. Thus, for example, in some longitudinal studies, serial sampling from the same patient [9] along several weeks (or years) is being used for the follow up of some intestinal pathologies, for studying the evolution of the gut microbiome after antibiotic treatment, or for colon cancer early detection [10] [11]. This need of sampling across time adds more complexity to metagenomics data storage and demands adapted algorithms to detect state variations across time as well as idiosyncratic commonalities of the microbiome of each individual [12]. In addition to the intra-individual sampling-time dependence, metagenomic clinical test results vary depending on the specific region of extraction of the clinical specimen. This local variability adds complexity to the analysis since different localizations (different tissues, different anatomical regions, healthy or tumor tissues) are required to have a sufficiently complete landscape of the human microbiome. Moreover, re-analysis of old samples using new tools and better reference databases might be also demanded from time to time.

Other disciplines such as astronomy or particle physics have faced the big data challenge before. A key difference is the existence of standards for data processing [7]; in metagenomics global standards for converting raw sequence data into processed data are not yet well defined, and there are shortcomings derived from the fact that most bioinformatics methodologies used for metagenomics data analysis were designed for scenarios very different from the current one. These are some of the aspects that have suffered crucial changes and advances with a direct impact in metagenomics data analysis:

1. **Sequence data:** the reads are larger, the sequencing depth and the number of samples of each project are considerably bigger. The first metagenomics studies were very local projects, while nowadays the most fruitful studies are done at a global level (international, continental, national). This kind of global studies has yielded the discovery of clinical biomarkers for diseases of the importance of cancer, obesity or inflammatory bowel diseases and has allowed exploring the biodiversity of varied earth environments.
2. **The genomics explosion:** its effect being felt in this case in the reference sequences. The immense amount of sequences available in public repositories demands new strategies for curation, update and storage of metagenomics reference databases: current models will (already) have problems to face the future avalanche of metagenomic sequence data.
3. **Cloud computing:** the appearance of new models for massive computation and storage such as the cloud-based platforms, or the widespread adoption of programming methodologies like functional programming, or, more speculatively, dependently typed programming. The new possibilities that these advances offer must have a direct impact in metagenomics data analysis.
4. **Open science:** the new social manner to do science, particularly so in genomics, brings its own set of requirements. Metagenomics evolves in a social and global scenario following a science democratization trend in which many small research groups from distant countries share a common big metagenomics project; this global cooperation demands systems allowing for reproducible data analysis, data interoperability, and tools and practices for asynchronous collaboration between different groups.

## Results

### Overview

Considering the current new metagenomics scenario and to tackle the challenges posed by metagenomics big data analysis outlined in the Introduction we have designed a new open source methodology for analyzing metagenomics data. It exploits the new possibilities that cloud computing offers to get a system robust, programmatically configurable, modular, distributed, flexible, scalable and traceable in which the biological databases of reference sequences can be easily updated and/or frequently substituted by new ones or by databases specifically designed for focused projects.

These are some of the more innovative MG7 features:

- Static reproducible specification of dependencies and behavior of the different components using *Statika* and *Datasets*
- Parallelization and distributed analysis based on AWS, with on-demand infrastructure as the basic paradigm
- Definition of complex workflows using *Loquat*, a composable system for scaling/parallelizing stateless computations especially designed for AWS
- A new approach to data analysis specification, management and specification based on working with it in exactly the same way as for a software project, together with the extensive use of compile-time structures and checks
- Modeling of the taxonomy tree using the new paradigm of graph databases (Bio4j). It facilitates the taxonomic assignment tasks and the calculation of the taxa abundance values considering the hierarchic structure of taxonomy tree (cumulative values)
- Exhaustive per-read taxonomic assignment using two complementary assignment algorithms Lowest Common Ancestor and Best BLAST Hit
- Using a new 16S database of reference sequences (16S-DB7) with a flexible and sustainable system of updating and project-driven customization

### Libraries and resources

In this section we describe the resources and libraries developed by the authors on top of which MG7 is built. All MG7 code is written in Scala, a hybrid object-functional programming language. Scala was chosen based on the possibility of using certain advanced programming styles, and Java interoperability, which let us build on the vast number of existing Java libraries; we take advantage of this when using Bio4j as an API for the NCBI taxonomy. It has support for type-level programming, type-dependent types (through type members) and singleton types, which permits a restricted form of dependent types where types can depend essentially on values determined at compile time (through their corresponding singleton types). Conversely, through implicits one can retrieve the value corresponding to a singleton type.

#### *Statika*: machine configuration and behavior

Statika is a Scala library developed by the first and last authors which serves as a way of defining and composing machine behaviors statically. The main component are **bundles**. Each bundle declares a sequence of computations (its behavior) which will be executed in an **environment**. A bundle can *depend* on other bundles, and when being executed by an environment, its DAG (Directed Acyclic Graph) of dependencies is linearized and run in sequence. In our use, bundles correspond to what an EC2 instance should do and an environment to an AMI (Amazon Machine Image) which prepares the basic configuration, downloads the Scala code and runs it.

#### *Datasets*: a mini-language for data

Datasets is a Scala library developed by the first and last authors with the goal of being a Scala-embedded mini-language for datasets and their locations. **Data** is represented as type-indexed fields: keys are modeled as singleton types, and values correspond to what could be called a denotation of the key: a value of type `Location` tagged with the key type. Then a **Dataset** is essentially a collection of data, which are guaranteed statically to be different through type-level predicates, making use of the value–type correspondence which can be established through singleton types and implicits. A dataset location is then just a list of locations formed by locations of each dataset key. All this is based on what could be described as an embedding in Scala of an extensible record system with concatenation on disjoint labels, in the spirit of [13] [14]. For that *Datasets* uses ohnosequences/cosas library.

Data keys can further have a reference to a **data type**, which, as the name hints at, can help in providing information about the type of data we are working with. For example, when declaring Illumina reads as a data, a data type containing information about the read length, insert size or end type (single or paired) is used.

A **location** can be, for example, an S3 object or a local file; by leaving the location type used to denote particular data free we can work with different “physical” representations, while keeping track of to which logical data they are a representation of. Thus, a process can generate locally a `.fastq` file representing the merged reads, while another can put it in S3 with the fact that they all correspond to the “same” merged reads is always present, as the data that those “physical” representations denote.

#### *Loquat*: Parallel data processing with AWS

Loquat is a library developed by the first, second and last authors designed for the execution of embarrassingly parallel tasks using S3, SQS and EC2 Amazon services.

A *loquat* executes a process with explicit input and output datasets (declared using the *Datasets* library described above). Workers (EC2 instances) read from an SQS queue the S3 locations for both input and output data; then they download the input to local files, and pass these file locations to the process to be executed. The output is then put in the corresponding S3 locations.

A manager instance is used to monitor workers, provide initial data to be put in the SQS queue and optionally release resources depending on a set of configurable conditions.

Both worker and manager instances are *Statika* bundles. The worker can declare any dependencies needed to perform its task: other tools, libraries, or data.

All configuration such as the number of workers or the instance types is declared statically, the specification of a loquat being ultimately a Scala object. Deploy and resource management methods make easy to use an existing loquat either as a library or from (for example) a Scala REPL.

The input and output (and their locations) being defined statically has several critical advantages. First, composing different loquats is easy and safe; just use the output types and locations of the first one as input for the second one. Second, data and their types help in not mixing different resources when implementing a process, while serving as a safe and convenient mechanism for writing generic processing tasks. For example, merging paired-end Illumina reads generically is easy as the data type includes the relevant information (insert size, read length, etc) to pass to a tool such as FLASh.

#### Type-safe eDSLs for BLAST and FLASh

We developed our own Scala-based type-safe eDSLs (embedded Domain Specific Languages) for FLASh [15] and BLAST [16] expressions and their execution.

In the case of BLAST we use a model where we can guarantee for each BLAST command expression at compile time that

- all required arguments are provided
- only valid options are provided
- correct types for each option value
- valid output record specification

Generic type-safe parsers returning a heterogeneous record of BLAST output fields are also available, together with output data defined using *Datasets* which have a reference to the exact BLAST command options which yielded that output. This lets us provide generic parsers for BLAST output which are guaranteed to be correct.

In the same spirit as for BLAST, we implemented a type-safe eDSL for FLASh expressions and their execution, supporting features equivalent to those outlined for the BLAST eDSL.

#### Bio4j and Graph Databases

Bio4j [17] is a data platform integrating data from different resources such as UniProt or GO in a graph data paradigm. In the assignment phase we use a subgraph containing the NCBI Taxonomy, wrapping in Scala its Java API in a tree algebraic data type.

#### 16S-DB7 Reference Database Construction

Our 16S-DB7 Reference Database is a curated subset of sequences from the NCBI nucleotide database **nt**. The sequences included were selected by similarity with the bacterial and archaeal reference sequences downloaded from the **RDP database** [18]. RDP unaligned sequences were used to capture new 16S RNA sequences from **nt** using BLAST similarity search strategies and then, performing additional curation steps to remove sequences with poor taxonomic assignments to taxonomic nodes close to the root of the taxonomy tree. All the nucleotide sequences included in **nt** database has a taxonomic assignment provided by the **Genbank** sequence submitter. NCBI provides a table (available at ftp://ftp.ncbi.nlm.nih.gov/pub/taxonomy/) to do the mapping of any Genbank Identifier (GI) to its Taxonomy Identifier (TaxID). Thus, we are based on a crowdsourced submitter-maintained taxonomic annotation system for reference sequences. It supposes a sustainable system able to face the expected number of reference sequences that will populate the public global nucleotide databases in the near future. Another advantageous point is that we are based on NCBI taxonomy, the *de-facto* standard taxonomic classification for biomolecular data [19]. NCBI taxonomy is, undoubtedly, the most used taxonomy all over the world and the most similar to the official taxonomies of each specific field. This is a crucial point because all the type-culture and tissue databanks follow this official taxonomical classification and, in addition, all the knowledge accumulated during last decades is referred to this taxonomy. In addition NCBI provides a direct connection between taxonomical formal names and the physical specimens that serve as exemplars for the species [20].

Certainly, if metagenomics results are easily integrated with the theoretical and experimental knowledge of each specific area, the impact of metagenomics will be higher than if it progresses as a disconnected research branch. Considering that metagenomics data interoperability, which is especially critical in clinical environments, requires a stable taxonomy to be used as reference, we decided to rely on the most widely used taxonomy: the NCBI taxonomy. In addition, the biggest global sequence database GenBank follows this taxonomy to register the origin of all their submitted sequences. Our 16S database building strategy allows the substitution of the 16S database by any other subset of **nt**, even by the complete **nt** database if it would be needed, for example, for analyzing shotgun metagenomics data. This possibility of changing the reference database provides flexibility to the system enabling it for easy updating and project-driven personalization.

### Workflow Description

The MG7 analysis workflow is summarized in Figure 1. The input files for MG7 are the FASTQ files resulting from a paired-end NGS sequencing experiment.

**Figure 1:**
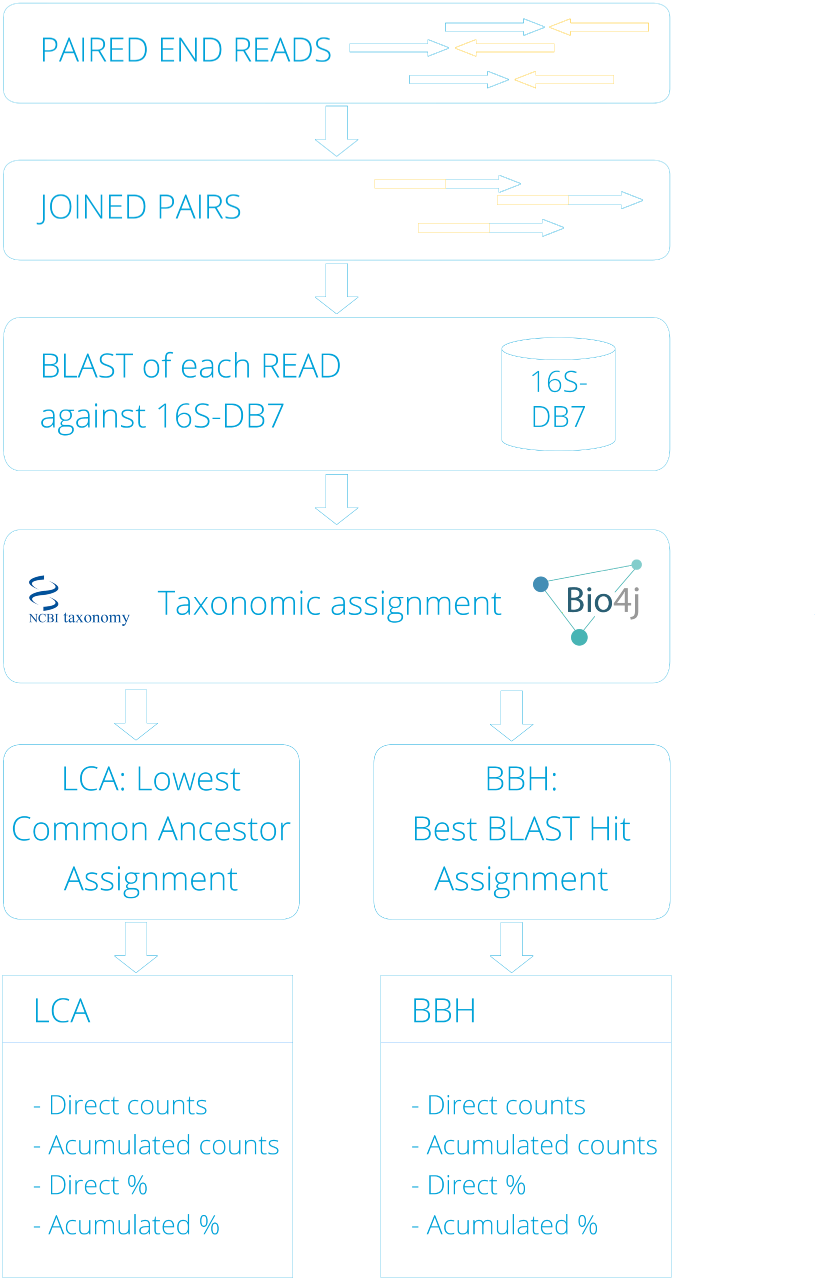
MG7 analysis workflow. The paired reads in fastq format are merged resulting in only one sequence per read pair. The next step is a parallelized BLASTN of every merged sequence against the 16S reference database 16S-DB7. Then, the mapping of the detected similar sequences in the database to the taxonomy node to which they belong is carried out. This is done using Bio4j that includes a module with all the NCBI taxonomy in a graph connected with the Gene Ontology, Uniprot, and RefSeq graphs. Then the taxonomic assignment is done for each sequence following two different approaches: LCA and BBH, and finally the abundances corresponding to direct and cumulative assignments for each node in percentage and absolute counts are provided for each assignment mode.

#### Joining reads of each pair using FLASh

In the first step the paired-end reads, designed with an insert size that yields pairs of reads with an overlapping region between them, are assembled using FLASh [15]. FLASh is designed to merge pairs of reads when the original DNA fragments are shorter than twice the length of reads. Thus, the sequence obtained after joining the 2 reads of each pair is larger and has better quality since the sequence at the ends of the reads is refined merging both ends in the assembly. To have a larger and improved sequence is crucial to do more precise the inference of the bacterial origin based on similarity with reference sequences.

#### Parallelized BLASTN of each read against the 16S-DB7

The second step is to search for similar 16S sequences in our 16S-DB7 database. The taxonomic assignment for each read is based on BLASTN of each read against the 16S database.Assignment based on direct similarity of each read one by one compared against a sufficiently wide database is considered in different reviews of metagenomics analysis methodologies [21] [22] as a very exhaustive method for assignment. Some methods of assignment compare the sequences only against the 16S genes from available complete bacterial genomes or avoid computational cost clustering or binning the sequences first, and then doing the assignments only for the representative sequence of each cluster. MG7 carries out an exhaustive comparison of all the reads under analysis and it does not applies any binning strategy. Every read is specifically compared with all the sequences of the 16S database. We select the best BLAST hits (10 hits by default) obtained for each read to do the taxonomic assignment.

#### Taxonomic Assignment Algorithms

All the reads are assigned under two different algorithms of assignment: i. Lowest Common Ancestor based taxonomic assignment (LCA) and ii. Best BLAST Hit based taxonomic assignment (BBH). Figure 2 displays schematically the LCA algorithm applied sensu stricto (left panel) and the called ‘in line’ exception (right panel) designed in order to gain specificity in the assignments in the cases in which the topology of the taxonomical nodes corresponding to the BLAST hits support sufficiently the assignment to the most specific taxon.

**Figure 2:**
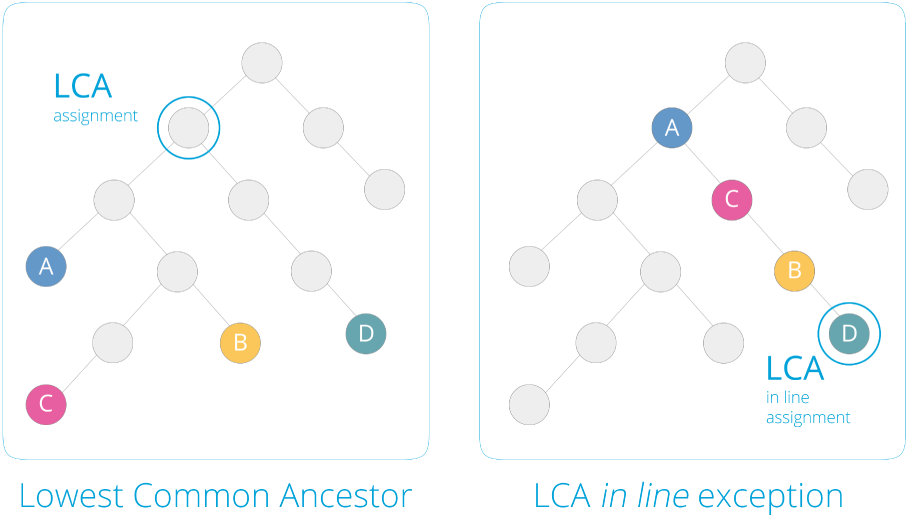
Lowest Common Ancestor algorithm for taxonomic assignment. The Left panel displays an example of the application of LCA algorithm in a *sensu stricto* mode. A, B, C and D represent taxonomy tree nodes with assigned reads. Right panel displays the *in line* mode of assignment which is an exception for the *sensu stricto* mode of application of LCA algorithm. The *in line* mode is used when all the nodes are located in a line without bifurcations. In that case the taxon assigned is the most specific (the most distant from the root).

**Lowest Common Ancestor based Taxonomic Assignment** For each read, first, we select the BEST BLAST HITs (by default 10 Hits) over a threshold of similarity. To evaluate similarity for this first filtering of hits we use the Expect value (by default *evalue ≤ e*^−15^) that describes the number of hits one can “expect” to see by chance when searching a database of a particular size. In a second filtering step we filtering those hits that are not sufficiently good comparing them with the best one. We select the best HSP (High Similarity Pair) per reference sequence and then choose the best HSP (that with lowest E-value) between all the selected ones. The bitscore of this best HSP (called S) is used as reference to filter the rest of HSPs. All the HSPs with bitscore below the product *pS* are filtered. p is a coefficient fixed by the user to define the bitscore required, e.g. if *p* = 0.9 and *S* = 700 the required bitscore threshold would be 630. Once we have the definitive HSPs selected, we obtain their corresponding taxonomic nodes using the taxonomic assignments that NCBI provides for all the nt database sequences. Now we have to analyze the topological distribution of these nodes in the taxonomy tree: i. If all the nodes forms a line in the taxonomy tree (are located in a not branched lineage to the tree root) we should choose the most specific taxID as the final assignment for that read. We call to this kind of assignment the ‘in line’ exception (see Figure 2 right panel). ii. If not, we should search for the *sensu stricto* Lowest Common Ancestor (LCA) of all the selected taxonomic nodes (See Figure 2 left panel). In this approach we decided to use the bitscore for evaluating the similarity because it is a value that increases when similarity is higher and depends a lot on the length of the HSP. Some reads could not find sequences with enough similarity in the database and then they would be classified as reads with no hits. Advanced metagenomics analysis approaches [23] have adopted LCA-based assignment algorithms because it provides fine and trusted taxonomical assignment.

**Best BLAST hit taxonomic assignment** We decided to maintain the simpler method of Best BLAST Hit (BBH) for taxonomic assignment because, in some cases, it can provide information about the sequences that adds information to that obtained using the LCA algorithm. With the LCA algorithm, when some reference sequences with BLAST alignments over the required thresholds map to a not sufficiently specific taxID, the read can be assigned to an unspecific taxon near to the root of the taxonomy tree. If the BBH reference sequence maps to more specific taxa, this method, in that case, gives us useful information.

#### Output for LCA and BBH assignments

MG7 provides independent results for the 2 different approaches, LCA and BBH. The output files include, for each taxonomy node (with some read assigned), abundance values for direct assignment and cumulative assignment. The abundances are provided in counts (absolute values) and in percentage normalized to the number of reads of each sample. Direct assignments are calculated counting reads specifically assigned to a taxonomic node, not including the reads assigned to the descendant nodes in the taxonomy tree. Cumulative assignments are calculated including the direct assignments and also the assignments of the descendant nodes. For each sample MG7 provides 8 kinds of abundance values: LCA direct counts, LCA cumu. counts, LCA direct %, LCA cumu. %, BBH direct counts, BBH cumu. counts, BBH direct % and BBH cumu. %.

#### Data analysis as a software project

The MG7 16S data analysis workflow is indeed a set of tasks, all of them based in *Loquat*. For each task, a set of inputs and outputs as well as configuration parameters must be statically defined. The user is also free to leave the reasonable defaults for configuration, needing only to define the input and output of the whole workflow. The definition of this configuration is Scala code and the way of starting an MG7 analysis is compiling the project code and launching it from the Scala interactive console.

Code compilation prior to launching any analysis assures that no AWS resources are launched if the analysis is not well-defined, avoiding expenses not leading to any analysis. Besides compile-time checks, runtime checks are made before launch to ensure existence of input data and availability of resources.

An MG7 analysis is then a Scala project where the user only needs to set certain variables at the code level (input, output and parameters), compile the code and run it. To facilitate the process of setting up the Scala project, a template with sensible defaults is provided.

In order to be able to exploit AWS infrastructure for the MG7 analysis, the user needs to set up an AWS account with certain IAM (Identity and Access Management) permission policies that will grant access to the resources used in the workflow.

### Availability

MG7 is open source, available at https://github.com/ohnosequences/mg7 under an AGPLv3 license.

## Discussion

We could summarize the most innovative ideas and developments in MG7:

1. Treating data analysis as a software project. This makes for radical improvements in *reproducibility*, *reuse*, *versioning*, *safety*, *automation* and *expressiveness*
2. Checking at compile-time: input and output data, their locations and type are expressible and checked at compile-time using *Datasets*
3. Management of dependencies and machine configurations using *Statika*
4. Automation of AWS cloud resources and processes, including distribution and parallelization through the use of *Loquat*
5. Taxonomic data and related operations are treated natively as what they are: graphs, through the use of *Bio4j*
6. MG7 provides a sustainable model for taxonomic assignment, appropriate to face the challenging amount of data that high throughput sequencing technologies generate

We will expand on each item in the following sections.

### A new approach to data analysis: data analysis as a software project and checking at compile-time

MG7 proposes to define and work with a particular data analysis task as a software project, using Scala. The idea is that *everything*: data description, their location, configuration parameters and the infrastructure used should be expressed as Scala code, and treated in the same way as any (well-managed) software project. This includes, among other things, using version control systems (git in our case), writing tests, making stable releases following semantic versioning or publishing artifacts to a repository.

What we see as key advantages of this approach (when coupled with compile-time specification and checking), are

- **Reproducibility** the same analysis can be run again with exactly the same configuration in a trivial way.
- **Versioning** as in any software project, there can be different versions, stable releases, etc.
- **Reuse** we can build standard configurations on top of this and reuse them for subsequent data analysis. A particular data analysis *task* can be used as a *library* in further analysis.
- **Decoupling** We can start working on the analysis specification, without any need for available data in a much easier way.
- **Documentation** We can take advantage of all the effort put into software documentation tools and practices, such as in our case Scaladoc or literate programming. As documentation, analysis processes and data specification live together in the files, it is much easier to keep coherence between them.
- **Expresiveness and safety** For example in our case we can choose only from valid illumina read types, and then build a default FLASh command based on that. The output locations, being declared statically, are also available for use in further analysis.

### Input and output data declaration

An important aspect of the MG7 workflow is the way it deals with data resources. All the data that is going to be used in the analysis or produced as an output is described as Scala code using rich types from the *Datasets* language. This allows the user to specify information about types of data, information that can then be utilized by tools analyzing this data. For example, we can specify that, for the first part of the MG7 workflow, running FLASh in parallel requires illumina paired end reads and produces joined reads.

On one hand, specification of the input data allows us to restrict its type and force users to be conscious about what they pass as an input. On the other hand, specification of the output data helps to build a workflow as a *composition* of several parts: we can ensure on the Scala code type level that the output of one component fits as an input for the next component. This is crucial as, obviously, the way a data analysis task works depends a lot on the particular structure of the data. For instance, in the MG7 workflow, using BLAST eDSL, we can precisely describe which format will have the output of the BLAST step, which information it will include, and then in the next step we can reuse this description to parse BLAST output and retrieve the part of the information needed for the taxonomy assignment analysis. Having the data structure described statically as Scala code allows us to be sure that we will not have parsing problems or other issues with incompatible data passed between workflow components.

All this does not compromise flexibility in how the user works with data in MG7: having static data declarations as a part of the configuration allows the user to reuse analysis components, or modify them according to particular needs. Besides that, an important advantage of the type-level control is the added protection from the execution (and deployment) of a wrongly configured analysis task, which may lead to significant costs in both time and money.

### Tools, data, dependencies and automated deployment

Bioinformatics software often has a complicated installation process and requires various dependencies with unclear versions. This makes the deployment of the bioinformatics tools an involved task and resolving it manually is not a solution in the context of cloud computations. To face this problem, one needs an automated system of managing tools and resources, which will allow an expressive way for describing dependencies between parts of a pipeline and provide a reproducible procedure of its deployment. We have developed *Statika* for this purpose and successfully used it in MG7.

Every external tool involved in the workflow is represented as a *Statika* bundle, which is essentially a Scala project describing the installation process of this tool and declaring dependencies on other bundles which will be installed prior to the considered tool itself. Describing relationships between bundles on the code level allows us to track the directed acyclic graph of their dependencies and linearize them to automatically install them sequentially in the right order. Meanwhile, describing the installation process on the code level allows the user to utilize the wide range of available Scala and Java APIs and tools, making installation a well-defined sequence of steps rather than an unreliable script, dependent on a certain environment. *Statika* offers an easy path towards making deployment an automated, reproducible process.

Besides bioinformatics tools like BLAST and FLASh, *Statika* bundles are used for wrapping data dependencies and all inner components of the system that require cloud deployment. In particular, all components of *Loquat* are bundles; the user can then define which components are needed for the parallel processing on each computation unit in an expressive way, declaring them as bundle dependencies of the loquat “worker” bundle. This modularization is also important for the matter of making components of the system reusable for different projects and liberating the user from most of the tasks related to their deployment.

### Parallel computations in the cloud

The MG7 workflow consists of certain steps, each of which performs some work in parallel, using the cloud infrastructure managed by *Loquat*. It is important to notice the horizontal scalability of this approach. Irrespectively of how much data needs to be processed, MG7 will handle it, by splitting data into chunks and performing the analysis on multiple computation units. The Amazon Elastic Compute Cloud (EC2) service provides a transparent way of managing computation infrastructure, called autoscaling groups. The User can set MG7 configuration parameters, adjusting for each task the amount and hardware characteristics of the EC2 instances they want to use for it. But it is important to note that, as each workflow step is not very resource demanding, it is not needed to hire EC2 instances with some advanced hardware. Instead, an average type will work and you can reduce execution time by simply scaling out the number of instances.

### Taxonomy and Bio4j

The hierarchic structure of the taxonomy of the living organisms is a tree, and, hence, is also a graph in which each node, with the exception of the root node, has a unique parent node. It led us to model the taxonomy tree as a graph using the graph database paradigm. Previously we developed Bio4j [17], a platform for the integration of semantically rich biological data using typed graph models. It integrates most publicly available data linked with sequences into a set of interdependent graphs to be used for bioinformatics analysis and especially for biological data. MG7 works based on the Bio4j taxonomy module. It opens the possibility to connect the taxonomic profiling data obtained with MG7 to all the biological knowledge associated to each taxon. Using the information available in Bio4j for all the proteins assigned to each taxon we are connected to all the functional data available in Uniprot related with it.

### Future developments

#### Shotgun metagenomics

It is certainly possible to adapt MG7 to work with shotgun metagenomics data. Simply changing the reference database to include whole genome sequence data could yield interesting results. This could also be refined by restricting reference sequences according to all sort of criteria, like biological function or taxonomy. Bio4j would be an invaluable tool here, thanks to its ability to express complex predicates on sequences using all the information linked with them (GO annotations, UniProt data, NCBI taxonomy, etc).

#### Comparing groups of samples

The comparison of the taxonomic profiles between different groups of samples is a need for many metagenomics studies. Tasks related with this group-based analysis, such as the extraction of the minimal tree with all the taxa with some direct or accumulated assignment, will be part of a new MG7 module, already in development.

#### Interactive visualizations based on Biographika

New visualization tools for metagenomics results are undoubtedly needed. Interactivity is a especially interesting feature for metagenomics data visualization, since the expert needs to explore the results in a knowledge-driven way. The majority of the available metagenomics data visualizations are static. We are working in the *Biographika* project [24], to provide interactive rich visualizations on the web for Bio4j data. The development of visualizations specific for MG7 is one of Biographika current goals. Biographika is based on D3.js, the de-facto standard JavaScript data visualization library, and is open source.

## Materials and Methods

### Amazon Web Services

MG7 uses the following Amazon Web Services:

- EC2 (Elastic Compute Cloud) autoscaling groups for launching and managing computation units
- S3 (Simple Storage Service) for storing input and output data
- SQS (Simple Queue Service) for communication between different components of the system
- SNS (Simple Notification Service) for e-mail notifications

These services are used through a Scala wrapper of the official AWS Java SDK v1.9.25: ohnosequences/aws-scalatools v0.13.2.

### Scala

MG7 itself and all the libraries used are written in Scala v2.11.

### Statika

MG7 uses ohnosequences/statika v2.0.0 for specifying the configuration and behavior of EC2 instances.

### Datasets

MG7 uses ohnosequences/datasets v0.2.0 for specifying input and output data, their type and their location.

### Loquat

MG7 uses ohnosequences/loquat v2.0.0 for the specification of data processing tasks and their execution using AWS resources.

### BLAST eDSL

MG7 uses ohnosequences/blast v0.2.0. The BLAST version used is v2.2.31+.

### FLASh eDSL

MG7 uses ohnosequences/flash v0.1.0. The FLASh version used is v1.2.11.

### Bio4j

MG7 uses bio4j/bio4j v0.12.0-RC3 and bio4j/bio4j-titan v0.4.0-RC2 as an API for the NCBI taxonomy.

## Disclosure/Conflict-of-Interest Statement

All authors work at the *Oh no sequences!* research group, part of Era7 Bioinformatics. Era7 offers metagenomics data analysis services based on MG7. MG7 is open source, available under the OSI-approved AGPLv3 license.

## Author Contributions

- **AA** developed *MG7*, *Statika*, *Datasets*, and *aws-scala-tools*; wrote the paper;
- **EK** developed *nispero* (a prototype for *Loquat* [25]) and *aws-scala-tools*.
- **MM** *MG7* workflow design; curation and design of the *16S-DB7* reference database; wrote the paper.
- **PPT** curation, design, data extraction code of the *16S-DB7* reference database.
- **EP** *MG7* workflow design; wrote the paper.
- **RT** *MG7* workflow design, assignment strategy; curation and design of the *16S-DB7* reference database; wrote the paper.
- **EPT** developed *MG7*, *Statika*, *Datasets*, *FLASh/BLAST eDSLs*; data analysis approach and design; wrote the paper.

All authors have read and approved the final manuscript.

## Acknowledgements

*Funding:* The two first authors are funded by INTERCROSSING (Grant 289974).

## References

[1] A. Oulas, C. Pavloudi, P. Polymenakou, G. A. Pavlopoulos, N. Papanikolaou, G. Kotoulas, C. Arvanitidis, and I. Iliopoulos Metagenomics: Tools and insights for analyzing next-generation sequencing data derived from biodiversity studies Bioinformatics and biology insights 9, 75. 2015.

[2] S. Bikel, A. Valdez-Lara, F. Cornejo-Granados, K. Rico, S. Canizales-Quinteros, X. Soberón, L. Del Pozo-Yauner, and A. Ochoa-Leyva Combining metagenomics, metatranscriptomics and viromics to explore novel microbial interactions: Towards a systems-level understanding of human microbiome Computational and structural biotechnology journal 13, 390–401. 2015.

[3] L. Ufarté, G. Potocki-Véronèse, and E. Laville Discovery of new protein families and functions: New challenges in functional metagenomics for biotechnologies and microbial ecology. Name: Frontiers in Microbiology 6, 563. 2015.

[4] L. M. Coughlan, P. D. Cotter, C. Hill, and A. Alvarez-Ordóñez Biotechnological applications of functional metagenomics in the food and pharmaceutical industries Frontiers in microbiology 6 2015.

[5] D. A. Cowan, J.-B. Ramond, T. P. Makhalanyane, and P. De Maayer Metagenomics of extreme environments Current opinion in microbiology 25, 97–102. 2015.

[6] R. Kodzius and T. Gojobori Marine metagenomics as a source for bioprospecting Marine genomics 2015.

[7] Z. D. Stephens, S. Y. Lee, F. Faghri, R. H. Campbell, C. Zhai, M. J. Efron, R. Iyer, M. C. Schatz, S. Sinha, and G. E. Robinson Big data: Astronomical or genomical? PLoS Biol 13 (7): e1002195. 2015.

[8] E. C. Hayden Genome researchers raise alarm over big data Nature 2015.

[9] K. Faust, L. Lahti, D. Gonze, W. M. de Vos, and J. Raes Metagenomics meets time series analysis: Unraveling microbial community dynamics Current opinion in microbiology 25, 56–66. 2015.

[10] G. Zeller, J. Tap, A. Y. Voigt, S. Sunagawa, J. R. Kultima, P. I. Costea, A. Amiot, J. Böhm, F. Brunetti, N. Habermann, et al. Potential of fecal microbiota for early-stage detection of colorectal cancer Molecular systems biology 10 (11): 766. 2014.

[11] W. S. Garrett Cancer and the microbiota Science 348 (6230): 80–86. 2015.

[12] E. A. Franzosa, K. Huang, J. F. Meadow, D. Gevers, K. P. Lemon, B. J. Bohannan, and C. Huttenhower Identifying personal microbiomes using metagenomic codes Proceedings of the National Academy of Sciences, 201423854. 2015.

[13] R. W. Harper and B. C. Pierce Extensible records without subsumption 1990.

[14] R. Harper and B. Pierce A record calculus based on symmetric concatenation in Proceedings of the 18th ACM SIGPLAN-SIGACT symposium on Principles of programming languages ACM 1991, 131–142.

[15] T. Magoč and S. L. Salzberg Flash: Fast length adjustment of short reads to improve genome assemblies Bioinformatics 27 (21): 2957–2963. 2011.

[16] C. Camacho, G. Coulouris, V. Avagyan, N. Ma, J. Papadopoulos, K. Bealer, and T. L. Madden Blast+: Architecture and applications BMC bioinformatics 10 (1): 421. 2009.

[17] P. Pareja-Tobes, R. Tobes, M. Manrique, E. Pareja, and E. Pareja-Tobes Bio4j: A high-performance cloud-enabled graph-based data platform BioRxiv, 016758. 2015.

[18] J. R. Cole, Q. Wang, J. A. Fish, B. Chai, D. M. McGarrell, Y. Sun, C. T. Brown, A. Porras-Alfaro, C. R. Kuske, and J. M. Tiedje Ribosomal database project: Data and tools for high throughput rrna analysis Nucleic acids research, gkt1244. 2013.

[19] G. R. Cochrane and M. Y. Galperin The 2010 nucleic acids research database issue and online database collection: A community of data resources Nucleic acids research 38 (suppl 1): D1–D4. 2010.

[20] S. Federhen Type material in the ncbi taxonomy database Nucleic acids research, gku1127. 2014.

[21] N. Segata, D. Boernigen, T. L. Tickle, X. C. Morgan, W. S. Garrett, and C. Huttenhower Computational meta’omics for microbial community studies Molecular systems biology 9 (1): 666. 2013.

[22] X. C. Morgan and C. Huttenhower Chapter 12: Human microbiome analysis PLoS Comput Biol 8 (12): e1002808. 2012.

[23] D. H. Huson and N. Weber Microbial community analysis using megan. Methods in enzymology 531, 465–485. 2012.

[24] P. P. Tobes, E. P. Tobes, M. Manrique, E. Pareja, and R. Tobes Biographika: Rich interactive data visualizations on the web for the research community BioRxiv, 021063. 2015.

[25] E. Kovach, A. Alekhin, M. Manrique, P. Pareja-Tobes, E. Pareja, R. Tobes, and E. Pareja-Tobes Nispero: A cloud-computing based scala tool specially suited for bioinformatics data processing. In IWBBIO 2014, 1414–1415.

